# Human commensal *Candida albicans* strains demonstrate substantial within-host diversity and retained pathogenic potential

**DOI:** 10.1101/2022.09.09.507247

**Authors:** Faith M Anderson, Noelle Visser, Kevin Amses, Andrea Hodgins-Davis, Alexandra M Weber, Katura M Metzner, Michael J McFadden, Ryan E Mills, Matthew J O’Meara, Timothy Y James, Teresa R O’Meara

**Author notes:** Corresponding author Address correspondence to: Teresa O’Meara, Ph.D Department of Microbiology and Immunology 1150 W. Medical Center Dr Medical Sciences Building II, Room 6751 Ann Arbor, MI, 48109, USA. Department of Biology, University of Louisville, Louisville, Kentucky, USA. Department of Botany and Plant Pathology, Oregon State University, Corvallis, OR, USA. authors contributed equally (Faith M Anderson, Noelle Visser).

## Abstract

*Candida albicans* is a frequent colonizer of human mucosal surfaces as well as an opportunistic pathogen. *C. albicans* is remarkably versatile in its ability to colonize diverse host sites with differences in oxygen and nutrient availability, pH, immune responses, and resident microbes, among other cues. It is unclear how the genetic background of a commensal colonizing population can influence the shift to pathogenicity. Therefore, we undertook an examination of commensal isolates from healthy donors with a goal of identifying site-specific phenotypic adaptation and genetic variation associated with these phenotypes. We demonstrate that healthy people are reservoirs for genotypically and phenotypically diverse *C. albicans* strains, and that this genetic diversity includes both SNVs and structural rearrangements. Using limited diversity exploitation, we identified a single nucleotide change in the uncharacterized *ZMS1* transcription factor that was sufficient to drive hyper invasion into agar. However, our commensal strains retained the capacity to cause disease in systemic models of infection, including outcompeting the SC5314 reference strain during systemic competition assays. This study provides a global view of commensal strain variation and within-host strain diversity of *C. albicans* and suggests that selection for commensalism in humans does not result in a fitness cost for invasive disease.

## INTRODUCTION

*Candida albicans* is a common colonizer of humans, with between 20-80% of the world’s population estimated to be asymptomatically colonized at any given time [1, 2], although this depends on many factors, including host health status and diet [3–5]. Colonization occurs at multiple body sites including the mouth, skin, GI tract, and vaginal tract [6]. These sites present a wide range of physiological stresses to colonizing fungi, including variation in pH, temperature, and oxygen levels, as well as nutrient limitation and host immune responses [6–9]. *C. albicans*-host interactions are generally commensal, but *C. albicans* can also act as an opportunistic pathogen, resulting in an estimated 400,000 serious bloodstream infections per year [10–12]. Additionally, *C. albicans* can cause more minor mucosal infections, including oral and vaginal thrush and skin infections [13]. As a consequence, *C. albicans* represents the 2nd most common human fungal pathogen and the most common source of healthcare-associated fungal infections [14].

While host immune status is known to be an important predictor of disease outcomes, whether genetic variation between *C. albicans* strains also contributes to differences in virulence remains an open question. In the model yeast *Saccharomyces cerevisiae,* there is extensive variability in genotype and phenotype among different isolates that has been linked to the ability of *S. cerevisiae* to adapt to a wide range of environmental conditions [15, 16]. Recent work has highlighted intra-species variation in important aspects of *C. albicans* biology [17–20]. There are currently 17 clades of *C. albicans,* which were initially defined through multilocus sequence typing (MLST) [21–23]. More recently, genome sequencing has resolved finer population structure in the group [24]. Genetic epidemiology studies suggest that the major clades of *C. albicans* may differ in how frequently they are isolated from bloodstream infections or asymptomatic colonization [25], with five clades accounting for the majority of clinical isolates [26–28]. These clinical isolates have demonstrated significant variation in murine models of systemic infection, biofilm formation, cell wall remodeling, secretion of toxins and proteolytic enzymes, and morphological plasticity [25,29–33]. However, our primary understanding of the genotype-phenotype relationship in *C. albicans* results from analyses of a relatively limited set of pathogenic clinical isolates and their laboratory derivatives, with the majority of the work performed in the SC5314 genetic background. Detailed analysis of the genetic determinants of biofilm regulation between five different strains, each representing a different clade of *C. albicans,* have highlighted that circuit diversification is widespread between strains [17], adding complexity to our understanding of *C. albicans* biology.

Recent work suggests that *C. albicans* experiences fitness tradeoffs between invasive and colonizing growth, with selection for commensal behavior during colonization [34–40]. In serial passage experiments or competitive fitness experiments in the gut, mutations in key transcription factors controlling hyphal formation, *EFG1* and *FLO8*, resulted in increased fitness in the gut and decreased fitness in systemic models of infection [34–39]. In oral candidiasis, trisomic strains have been identified with a commensal phenotype [41]. Additionally, the 529L strain of *C. albicans* causes less damage and inflammation during oropharyngeal candidiasis [42], and persists at a higher fungal burden in both the mouth and the gut [20]. These potentially divergent selection pressures imply that commensal *C. albicans* strains can differ from isolates that cause invasive disease, but the genetic determinants underlying this difference have not been defined.

Here, we demonstrate that healthy people are reservoirs for genotypically and phenotypically diverse *C. albicans* strains that retain their capacity to cause disease. We obtained representative isolates from healthy undergraduate student donors, including both oral and fecal isolates. These isolates included representatives from 8 clades of *C. albicans,* including instances of colonization by strains from multiple clades within a single individual, highlighting the within-host diversity of the colonizing fungal population. Synthetic long-read sequencing coupled with CHEF gel analysis revealed large structural variation between strains, consistent with early karyotyping work in the field [43]. Phenotyping of the strains revealed extensive variation in growth, stress response, biofilm formation, and interaction with macrophages, but no separation of strains by sample origin site. Instead, we identified shared structural variations and phenotypes that were shared between strains from the same phylogenetic clade, suggesting that underlying genetic background may be more predictive of certain phenotypes than selection pressure at a specific sample site. We also discovered and mapped specific nucleotide variants to the ability of strains to invade agar, identifying a new role for *ZMS1* transcription factor. Notably, the majority of the strains showed increased competitive fitness in invertebrate models of systemic candidiasis compared with the reference SC5314 strain and retained their capacity to cause disease during monotypic infections. Together, these data suggest that the selective pressures experienced by *C. albicans* during commensal colonization do not necessarily result in decreased pathogenic potential.

## RESULTS

### Phenotypic Characterization of commensal *C. albicans* strains

We sourced *C. albicans* from oral and fecal samples from undergraduate student donors, using these as a representative sample of colonizing *C. albicans* strains from healthy individuals. These new isolates complement previous work focusing on bloodstream or mucosal infection isolates [18]. In this population of students, 16% (16/98) were positive for oral *Candida* colonization and 12% (12/98) were positive for fecal colonization (Fig 1A). From each host and site, we collected every individual colony present on the BD ChromAgar plates and confirmed species identity through ITS amplicon sequencing. Overall, we obtained 910 *C. albicans* isolates (fecal: n = 84 colonies, oral: n = 826 colonies), with oral samples giving rise to more individual colonies per donor (Fig 1B).

**Fig 1.**
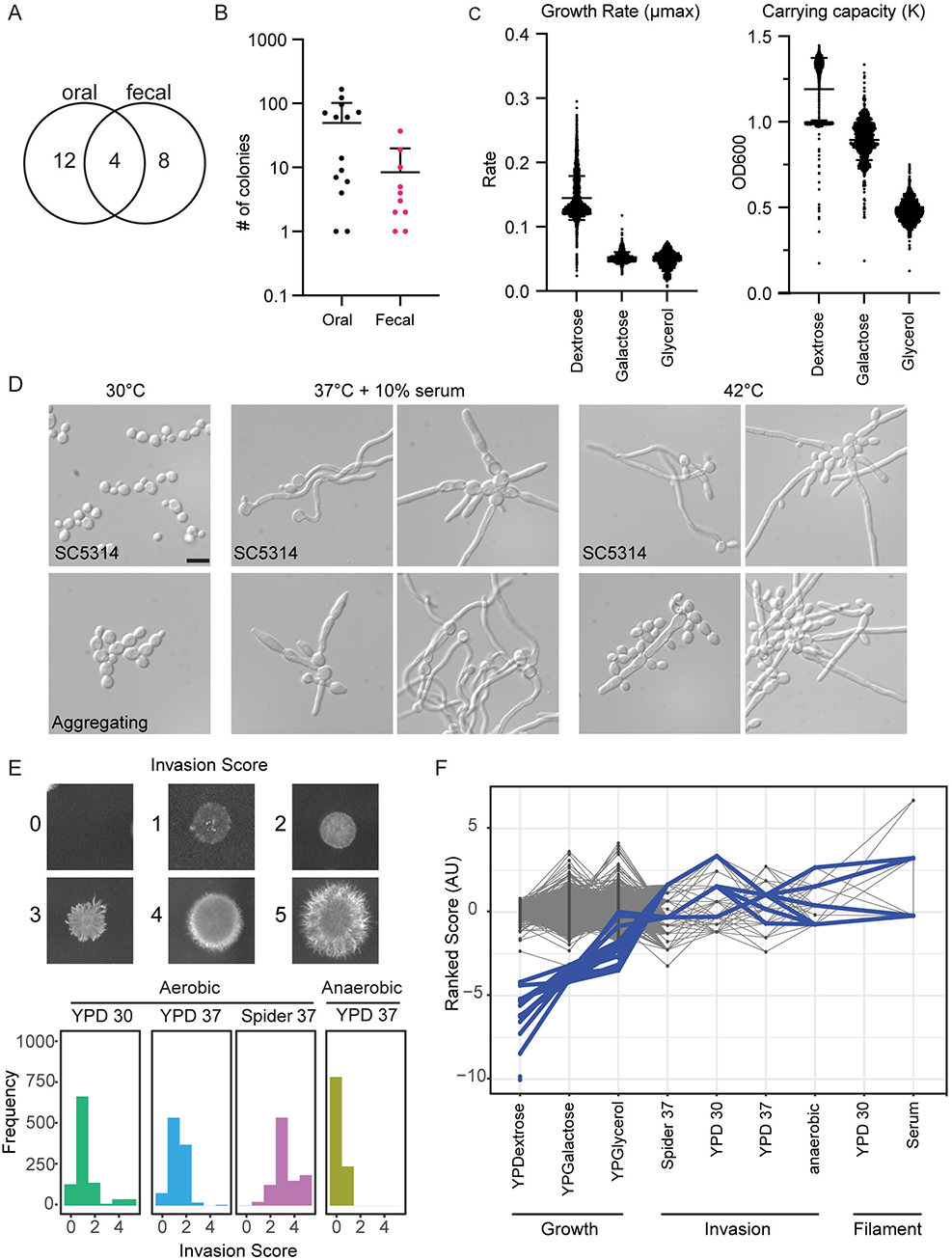
Characterization of isolates from healthy donors reveals extensive phenotypic heterogeneity. A) The number of healthy donors from whom *C. albicans* colonies were obtained by isolation site with 4 donors exhibiting both oral and fecal colonization. B) Number of *C. albicans* colonies isolated per donor from each sample site. Error bars represent mean and SD in number of colonies from each positive individual donor. C) Strains varied in maximum growth rate and carrying capacity in response to different carbon sources. Growth curves were performed on each isolate under three carbon sources at 30°C for 24 hours in biological duplicate. Rate and carrying capacity were determined using the GrowthcurveR analysis package. D) All strains retained the capacity to filament in liquid inducing cues, but some demonstrated altered morphology and aggregation. Strains were incubated in the indicated conditions and imaged at 20X magnification. Scale = 10 μM. E) Strains varied in their capacity to invade into solid agar. Colonies were incubated on the indicated conditions for 5 days before gentle washing and imaging for invasion. F) Parallel coordinate plot. Each strain was ranked for each growth condition, and the relationships between phenotypes are indicated with the lines. Blue highlights the slow growing strains.

We then performed growth assays on all 910 isolates and the SC5314 reference strain in rich medium (yeast peptone) with dextrose, galactose, or glycerol as the carbon source. We observed unimodal distributions with a long tail of slow-growing strains for both exponential growth (*µ_max_*) and saturating density (*K*) in rich media (Fig 1C). The five slowest-growing strains originated from five different hosts and showed significantly decreased growth rate and saturating density relative to other isolates from the same host. Although the five slowest-growing isolates were all obtained from oral samples, we did not observe a significant difference in growth rate between oral and fecal samples (S1A Fig), and overall, the growth rates between the different carbon sources were correlated (S1B Fig), as slow growth on one carbon source was predictive of slow growth on other carbon sources.

As filamentation has been tightly associated with virulence, and because previous work in murine models has suggested that gut adapted strains may lose their ability to filament [20,34,36], we examined each isolate in the collection for their ability to form hyphae using 10% serum and febrile temperatures as two inducing cues. Additionally, we examined the morphology of each strain under the non-inducing condition of YPD at 30°C to identify constitutive filamentation, as has been observed from isolates collected from sputum samples from cystic fibrosis patients [44]. We observed that while none of the strains were constitutively filamentous, several strains aggregated at YPD 30°C. All strains were able to filament in response to the inducing conditions of serum and high temperature, albeit with some variation in the shape and aggregation of the cells (Fig 1D). This may be consistent with the hypothesis that interaction with bacteria in the gut maintains selection for the hyphal program [34, 45]. Notably, the aggregative isolates were more likely to grow slowly and reach a lower carrying capacity. The universal ability of commensal strains isolated from healthy humans to filament stands in contrast to the observation that *C. albicans* evolves a yeast-locked phenotype in germ-free or antibiotic-treated mouse models, suggesting that selection in these models may not recapitulate important features of *C. albicans*-human interactions.

Filamentation programs under solid and liquid growth can involve distinct genetic programs [46], and the human host can present a variety of substrates for *C. albicans* to utilize. Therefore, we also tested each isolate for its colony morphology and capacity to invade into a set of solid agar media, including YPD agar at 30°C as the baseline condition, Spider agar at 37°C as a strong hyphae-inducing condition, YPD agar at 37°C as an intermediate inducing condition, and YPD agar at 37°C anaerobic conditions [46]. After several days of growth in each condition, colonies were gently rinsed from the plate and were given an invasion score from 0-5, with representative images of each score provided in (Fig 1E). As expected for a strong inducing cue, the highest degree of invasive growth was observed on Spider agar at 37°C [46]. Interestingly, although all strains showed the ability to form hyphae under liquid culture conditions, many strains failed to invade into solid agar, giving scores of 0 or 1 under multiple conditions (Fig 1E). Overall, there was a trend for the oral strains to invade more on Spider agar (S1C Fig). We also observed substantial phenotypic heterogeneity among strains isolated from the same host, including in strains isolated from the same site, consistent with each individual being colonized by multiple, phenotypically-distinct, strains of *C. albicans*.

To examine the relationship between the strains and the observed phenotypes, we generated a parallel coordinate plot by ranking each strain for each phenotype (Fig 1F). From these data, we could observe that a subset of the slow growing strains in YPD (blue) exhibited increased invasion on solid agar surfaces (Fig 1F), suggesting that slow growth in liquid media does not necessarily correlate with defects in other growth conditions. Upon closer inspection, several of these slow growing strains originated from donor 882. This subset of strains showed hyper invasion (score of 5) at the baseline condition of YPD agar at 30°C. Interestingly, although these strains retained their hyper invasive phenotype under the strong inducing cue of Spider agar at 37°C, they exhibited only moderate invasion (average score of 2) in the YPD agar at 37°C condition. Together, these experiments demonstrate a large variation in phenotypes between commensal isolates, even among traits, such as filamentation and invasion, that are often correlated with virulence.

### Genomic Variability

Due to the extensive variation in observed phenotypes among the isolates, we next wanted to characterize the genomic variability in these strains, at both a sequence and structural variation level. To carry out these analyses, we selected a set of 45 commensal isolates, hereafter referred to as the ‘condensed set’. Isolates were chosen for inclusion in the condensed set where we had matched pairs from oral and fecal isolates from the same donor or isolates from the same donor that exhibited multiple phenotypes based on growth, filamentation, or invasion. For these analyses, we compared each isolate to the SC5314 reference strain. We also included the human isolate, CHN1, to represent an alternate clinical isolate that has been previously phenotypically and genotypically characterized [47].

Previous descriptions of population genomic variation in *C. albicans* have largely relied on short read sequencing approaches, which are limited in their ability to resolve structural variation between strains. Therefore, we performed whole genome sequencing on all 45 commensal strains and the SC5314 and CHN1 reference strains using the Transposase Enzyme Linked Long-read Sequencing (TELL-Seq) method for library preparation [48]. This approach uses barcode linked-reads to produce synthetic long reads with Illumina quality sequence, thus allowing us to capture both SNVs and structural variants. The TELL-Seq library was then sequenced using an Illumina NovaSeq, resulting in a read coverage of approximately 150X for each sample.

To identify single nucleotide variants (SNVs) and compare our strains to the existing set of sequenced *C. albicans* isolates, we collected 388 previously published *C. albicans* genomes and mapped them, and our 45 newly-generated genomes, to the SC5314 reference genome with BWA-MEM [49]. We called variants using GATK HaplotypeCaller and after filtering, we obtained final set of 112,136 high quality SNVs across 431 remaining samples. Of these, 90,675 (80.86%) were represented in at least one member of our set of newly sequenced strains. The population diversity captured in this analysis was consistent with the largest previous analysis of *C. albicans* genomic variation, which called 589,255 SNPs from 182 genomes [24]. Although our final SNV set was significantly smaller than that identified in past work, our filtering criteria were significantly more stringent and robust for inferring population level patterns.

We then wanted to place our newly sequenced isolates in the phylogenetic context of previous work on *C. albicans* strains [18,24,50]. To remove redundancy and focus on natural *C. albicans* diversification, we removed samples corresponding to resequenced strains (e.g., multiple SC5314 samples present in full data set) and those sequenced as part of experimental evolution studies (i.e., [34,51,52]). Following removal of these samples, we were left with 324 sample SNV profiles, which were then used to cluster the samples into a dendrogram of relationships. Despite the reduced size of our dataset, our SNV-based clustering recovers all major accepted clades of *C. albicans* (Fig 2) [21–25]. The fact that >80% of the 112,136 high-quality SNVs we identified are represented in the genome sequences of the 45 new *C. albicans* isolates we sequenced, in addition to their clustering with 8 major clades, asserts the high degree of diversity captured in our study of commensal *C. albicans* strains actively colonizing humans. In line with past work and underpinning the validity of our SNV-based clustering, our isolates are not represented in clades of *C. albicans* known to exhibit a high degree of geographic specificity (e.g., Clade 13) (Fig 2) [24].

**Fig 2.**
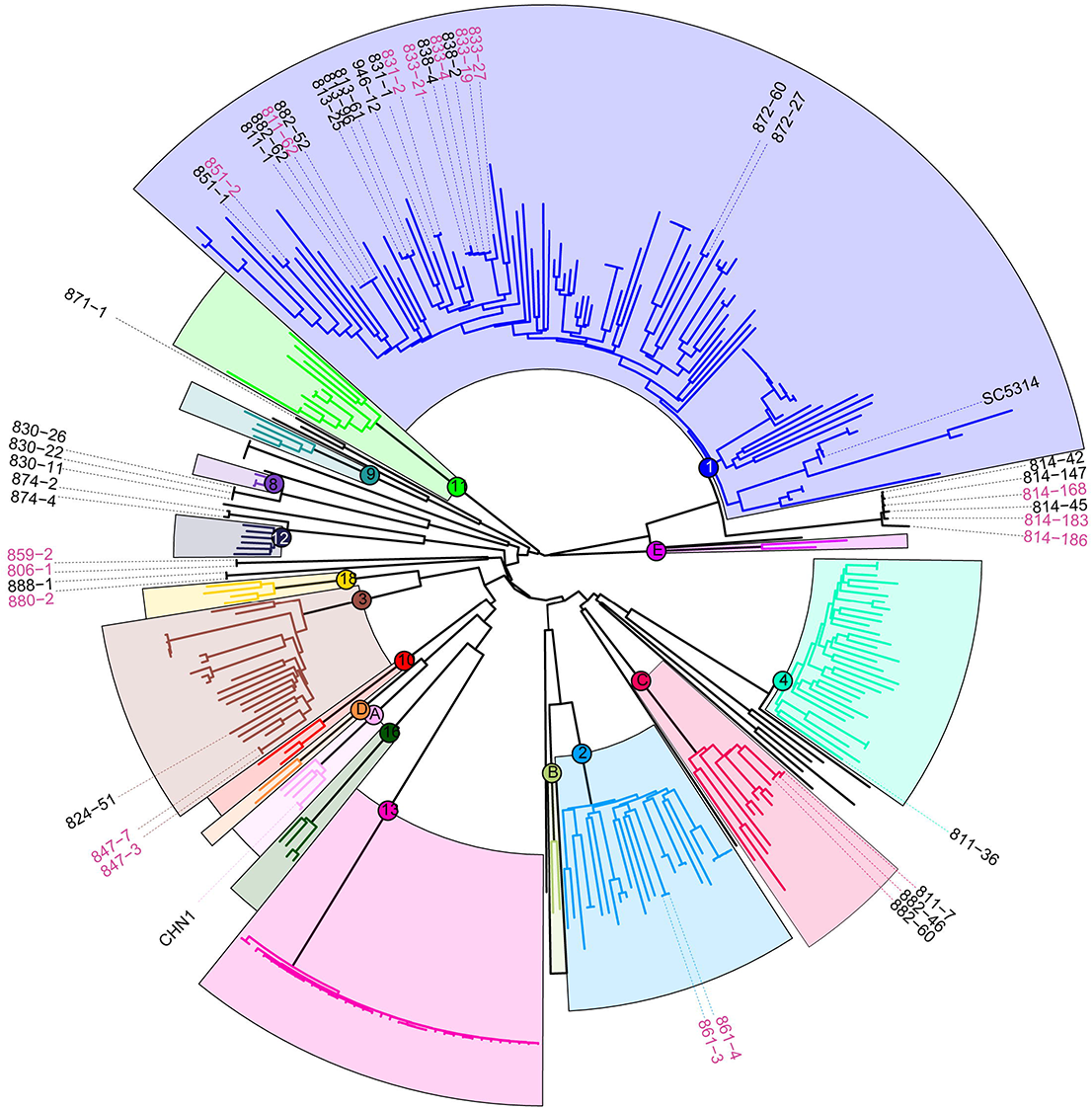
The newly sequenced *C. albicans* commensal isolates include representatives from multiple clades. A) A maximum likelihood tree showing the phylogenetic relationships between the 324 isolates analyzed via (neighbor joining). Previous clusters from [24] are highlighted with colored boxes. New isolates were colored based on the nearest defined cluster.

The majority of the new isolates (26/45) belonged to Clade 1, of which the reference strain SC5314 is also a member. We identified cases where isolates from the same donor clustered tightly together, such as donor 814, whose 6 strains included in the condensed set clustered in Clade 1. Consistent with previous reports on microevolution in the host [52], we observed primarily SNVs and short-tract loss of heterozygosity between these six isolates, perhaps consistent with clonal expansion and diversification during colonization. In contrast, we also identified donors with colonizing strains from multiple clades, such as donor 882, whose 4 strains came from Clade 1 and Clade C, or donor 811, whose 4 strains came from Clade 1, Clade C, and Clade 4 (Fig 2). Interestingly, some isolates from multiple donors clustered within one another, such as those from donors 838 and 833, likely indicating transmission between individuals. Together, this suggests that variation in an individual’s colonizing *C. albicans* strains can come from both within-host diversification and between-host transmission.

To characterize genomic variation at a structural level, we performed pulsed-field gel electrophoresis to separate the chromosomes of our condensed set of commensal *C. albicans* isolates, including SC5314 as a reference (Fig 3A, S2A Fig). The commensal *C. albicans* strains show between 7 and 10 chromosome bands, ranging from ∼0.8 MB to ∼3.2 MB in size [53]. We observed many size differences in chromosomes, especially among the smaller chromosomes (corresponding with Chr 5,6, and 7 in SC5314), but we also observed size variation in large chromosomes at approximately ∼ 1 MB, ∼1.5 MB, and ∼2 MB, potentially indicating variation in Chrs 2,4, and R.

**Fig 3.**
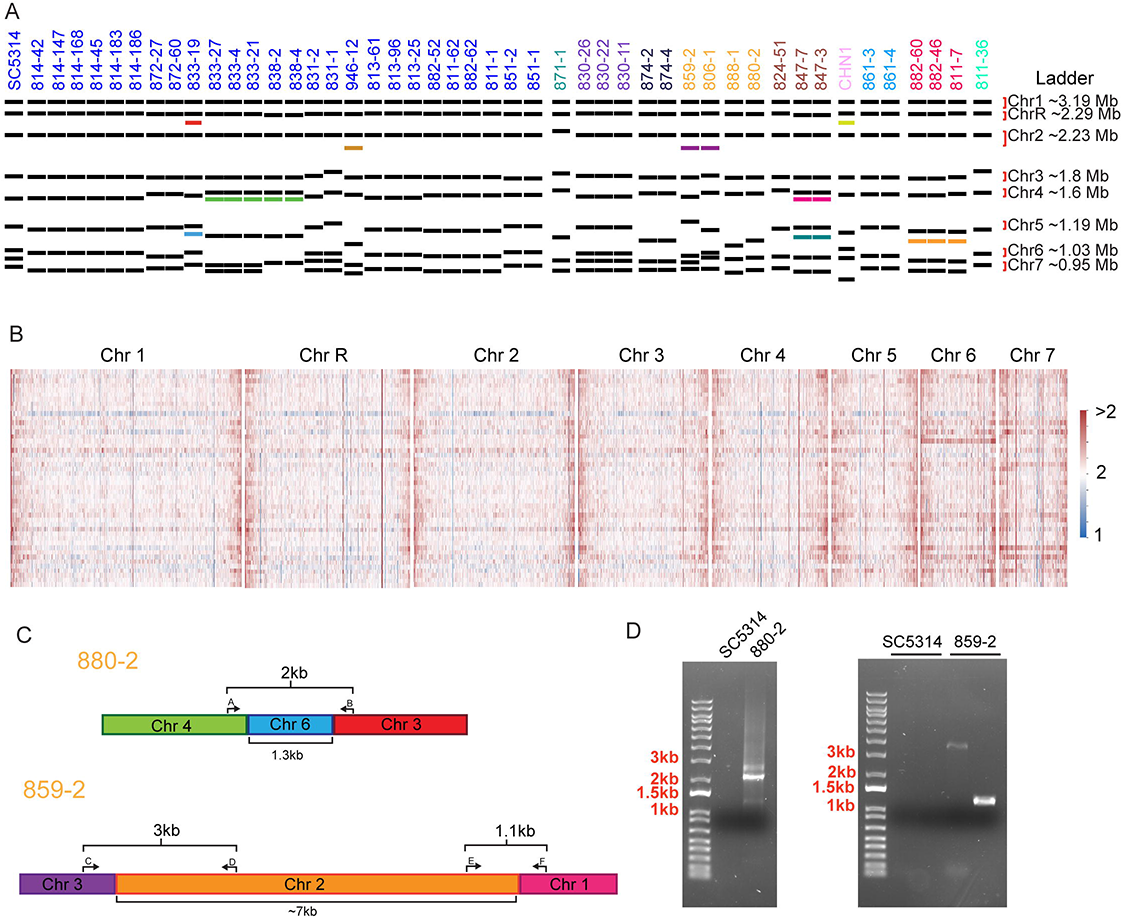
*C. albicans* commensal isolates display extensive structural variation. A) CHEF karyotyping gels show alterations in chromosome size and number between *C. albicans* isolates. Novel chromosomes are indicated with the colored bands. Gel images in S2 Fig. B) Heatmaps of read coverage across the condensed set of isolates for each chromosome. Each column represents a 500 bp bin of the reference genome and each row is an isolate from the condensed set. Values greater than one (red) suggest potential duplications while values less than one (blue) suggest potential deletions. C) Isolates 880-2 and 859-2 chromosomal fusion events identified & chosen for validation. D) PCR validation of fusion events in isolates 880-2 and 859-2. Bands corresponding with fusion events were not present in the SC5314 reference strain.

When comparing the chromosomal structural variations to our SNV tree, we identified several unique patterns, and observed that a phylogenetic ordering of the isolates did not encompass the structural variation. For example, a subset of isolates within Clade 1 from donors 833 and 838 all contained an extra chromosome below Chr 4, with the exception of isolate 833-19, despite being within the same phylogenetic cluster based on SNV analyses (Fig 3A). We also observed clade-level variation in karyotypes: Isolate 871-1 was the only strain in the condensed set from Clade 9, and this strain showed a unique pattern with different banding patterns at Chrs 2, 3, and 4. The 3 strains from Clade C, which originated from donors 882 and 811, all contained a chromosome band between Chrs 5 and 6. Finally, the two closely related strains, 859-2 and 806-1, both contained a chromosome band between Chrs 2 and 3 (Fig 3A).

An advantage of our TELL-seq based platform was the opportunity to resolve the differences between the SNV-level analysis and the structural variants that we identified through the CHEF gels. We used the synthetic long-reads to assemble contigs for each strain, using Universal Sequencing’s TELL-seq pipelines: Tell-Read and Tell-Link. We could observe variation in copy number across the chromosomes (Fig 3B), as well as potential inversions, duplications, or deletions (S2B Fig). Across our isolates, we observed substantial copy number variation, with a trend towards increased copy number. We noted that the isolates globally exhibited copy number expansion in the telomeric regions, and that copy number increases were more common on smaller chromosomes. Overall, the considerable copy number variation among our commensal isolates is in line with past work suggesting that host environmental pressures induce changes in genome size and that increased ploidy enhances fitness within the host [54, 55].

Notably, we were able to identify structural rearrangements and putative chromosome fusions, and used PCR to test for the presence of the fusion event. In strain 880-2, we observed a fusion event between chromosomes 3 and 4, connected by a 1.3 kb intervening sequence (Fig 3C). This intervening sequence had 94% sequence identity to an intergenic sub-telomeric region of chromosome 6. To determine whether this was a true event or a sequencing artifact, we designed primers to span the junction and performed PCR to amplify the fusion (Fig 3D). Using this approach, we observed that in strain 880-2, there is a bona fide structural rearrangement that links chromosomes 3 and 4. In strain 859-2, we observed a fusion event between chromosomes 1 and 3, connected by an approximately 7kb intervening sequence with no obvious sequence identity to the SC5314 reference strain, but instead had 99.76% sequence identity to a region on chromosome 2 from *C. albicans* strain TIMM 1768 (Fig 3C). We were again able to use PCR to span both junctions observed the presence of the fusion between chromosomes 1, 2, and 3 (Fig 3D). This fusion event may correspond to the additional chromosomal band between chromosomes 2 and 3 that we observed in the karyotype for this strain. Importantly, neither of these fusion events were present in the SC5314 reference strain, indicating that the fusions were unique to the specific isolate (Fig 3D). These structural variations were not captured in the SNV analysis, and may play important roles in gene regulation or phenotypic variation between the strains.

Together, our sequence-level and structural-level genomic analysis of commensal *C. albicans* isolates indicate that healthy individuals can harbor multiple strains of *C. albicans* from different clades. Furthermore, our structural analysis shows substantial heterogeneity in number and size of chromosomes among commensal isolates, which suggests that even within similar strains, there is potential for structural variation.

### Deep Phenotyping of Commensal Isolates

The set of isolates for sequencing were initially chosen based on variation in growth rate in rich medium and alterations in invasion into agar. However, we hypothesized that we may identify specific adaptations in *C. albicans* strains isolated from different sites that allow for colonizing different host microenvironments. The host sites commonly colonized by *C. albicans* vary dramatically in environmental cues, such as nutrient availability, pH, immune responses, and resident microbiomes. Additionally, we hypothesized we may identify phenotypes associated with specific *C. albicans* clades, as we were able to identify structural variants shared between closely related isolates. To test this, we performed a set of growth analyses under multiple environmental conditions, including pH stresses, nutrient limitation, cell wall stressors, and antifungal drugs (Fig 4A). These analyses produced a dense array of quantitative phenotypic information for each strain.

**Fig 4.**
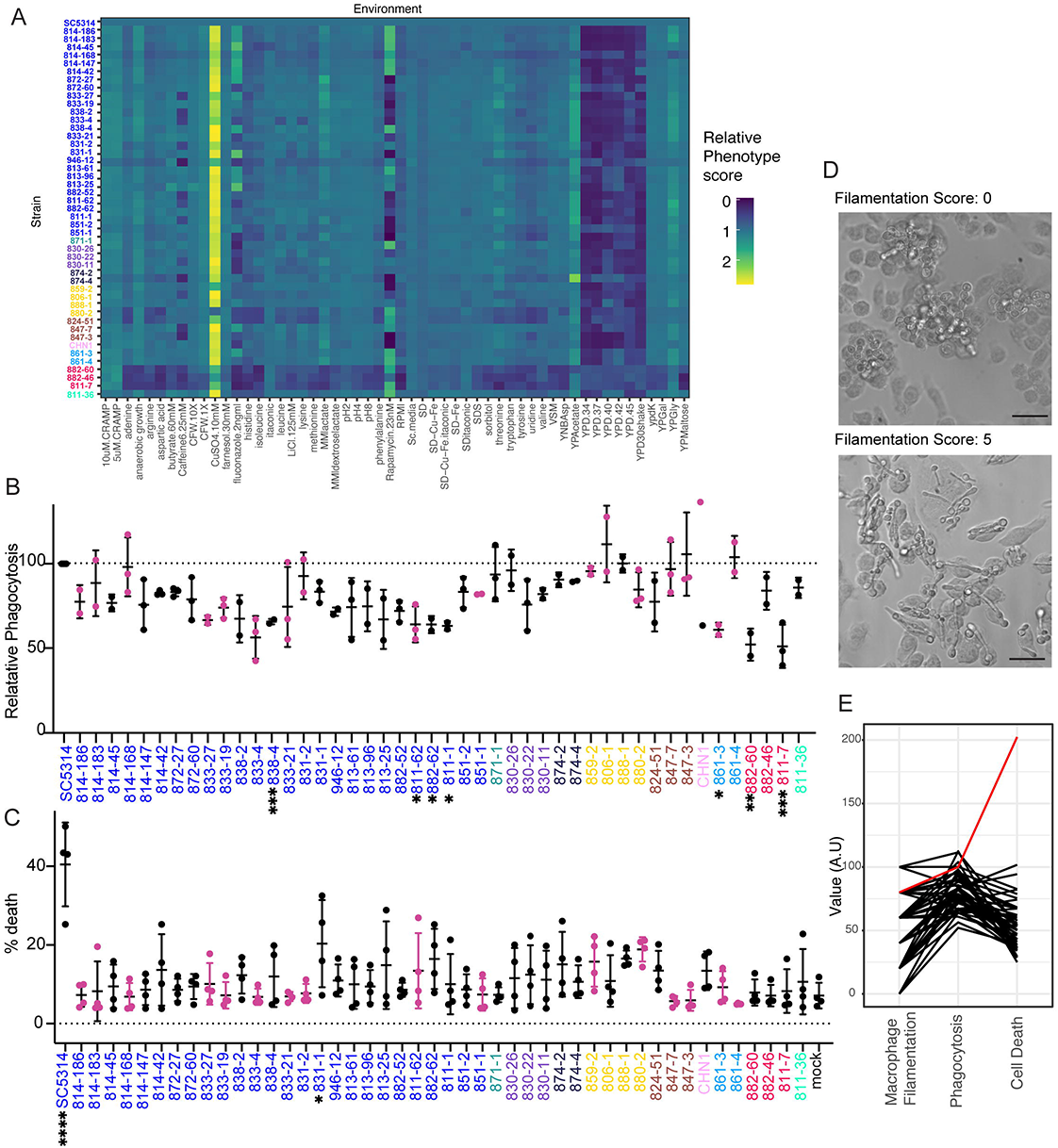
Deep phenotyping reveals heterogeneity in *in vitro* and host response phenotypes. A) Growth curve analysis under multiple environmental conditions. Carrying capacity (K) was normalized to SC5314, and the fold-change plotted by heatmap. Aggregating strains (882-60, 882-46, and 811-7) demonstrate a consistently lower carrying capacity. B) Relative macrophage phagocytosis rates of commensal isolates to reference strain SC5314. C) Macrophage cell death rates. D) Representative images of isolates following 4 hour macrophage infection. Representative filamentation score of 0 (left). Representative filamentation score of 5 (right). 20x magnification. Scale = 50 μM. E) Parallel coordinate plot ranking isolates for macrophage filamentation scoring, phagocytosis rates, and cell death rates. SC5314 reference strain is indicated in red. For phagocytosis rates, significant differences from the SC5314 reference strain were determined by one-way ANOVA, with Dunnett’s multiple correction testing. For cell death, significant differences from the mock condition were determined by one-way ANOVA, with Dunnett’s multiple correction testing. Asterisks indicate P < 0.05 (*), P < 0.005 (**), P < 0.001 (***) and P < 0.0001 (****).

From these data, we identified 3 strains, 882-60, 882-46, and 811-7, that consistently grew more poorly than the wildtype under multiple conditions; these strains were those that exhibited aggregation at 30°C and slow growth in rich media conditions. These strains all belonged to Clade C and were closely related, despite arising from two donors (Fig 4A). Growth rates in the nutrient limitation conditions were generally correlated with each other. However, we did not observe a correlation between body site and growth rate, even in response to cues that would appear to be specific for a particular body site, such as anaerobic growth. Across the commensal isolates, we noted the most variation in growth in response to caffeine and the antifungals fluconazole and rapamycin. In addition to growth, we measured each of the strains for their ability to form biofilms on plastic surfaces [56]. Although we observed variation between the strains, there was no correlation between isolation site or clade with the propensity of isolates to form biofilms (S3 Fig).

Our dense array of phenotypic data across 45 *C. albicans* isolates and 8 clades reveal that commensal isolates largely retain the plasticity to grow efficiently under diverse environmental cues, even those not immediately relevant to their colonizing site, as we did not observe growth enrichment in cues specific to isolation sites.

A major stress condition and environmental factor impacting *C. albicans* in the host is the immune response. Therefore, we moved from pure growth assays to measuring host-microbe interactions, using macrophages as representative phagocytes. We first hypothesized that the oral strains may show decreased recognition by macrophages, as persistent oral isolates were recently shown to result in reduced immune recognition and inflammation in both an OPC model of infection and in cell culture [57]. We tested this by measuring phagocytosis of each strain by immortalized bone-marrow derived macrophages and determining the ratio of internalized to external cells by differential staining and microscopy (Fig 4B) [58]. Although most isolates were not significantly different from the SC5314 reference, the isolates generally had a lower phagocytic rate than SC5314. Additionally, there was no correlation between sample origin site or clade with phagocytosis rate.

As phagocytosis was not a major differentiating factor between strains, we then wanted to examine whether the strains would induce different levels of macrophage cell death. We primed bone-marrow derived macrophage for 2 hours with LPS before infecting with each of our isolates for 4 hours. Following infection, we stained the cells with propidium iodide (PI) as a measure of cell death (Fig 4C). On average, the commensal isolates induced between 5% and 20% cell death, which was significantly lower than the reference strain, SC5314, which induced an average of 40% cell death. Several strains, including the 3 aggregating strains, 882-60, 882-46, and 811-7, were not significantly different than the mock condition. Other than SC5314, only one isolate, 831-1, was significantly different (p = 0.0411) from the mock condition.

Recently, we showed that *C. albicans* mutants that filament in serum are not always filamentous within macrophages [59]. As filamentation is linked, but not required, for inflammasome activation within host phagocytes [59–61], and clinical isolates show variability in induction of host inflammatory responses [62, 63], we examined the morphology of the commensal isolates after incubation for four hours with macrophages. We observed considerable variation in the extent of filamentation among the natural isolates (Fig 4D), including strains that completely failed to filament and those that filamented more than the SC5314 reference strain. Notably, the extent of filamentation did not correlate with colony morphology or invasion on agar, with many strains showing invasion into agar but no filamentation inside the macrophage, and vice versa (S4 Fig). Additionally, oral and fecal isolates both demonstrated defects in filamentation in macrophages, and filamentation in macrophages was not predictive of the phagocytic rate or cell death rate (Fig 4E).

Using individual phenotypic measures, we were unable to identify associations between strains based on body site or donor. However, it is possible that the combined phenotypic and genotypic profile would identify clusters of strains with similar distinct phenotypes or reveal connections between isolates. Therefore, we turned to uniform manifold approximation projection (UMAP) embedding, which will plot strains with similar phenotypes closer together and strains with dissimilar phenotypes farther apart based on the cosign metric. We observed three major clusters, but they did not segregate by isolation site, clade, or participant (S5 Fig). In sum, all of the commensal isolates showed extensive phenotypic variation, but this was not dependent on the body site or participant from which they were collected.

### Limited Diversity Exploitation

Genome-wide association studies have been a powerful tool for identifying the genetic basis of variation in phenotypes of interest in humans and other recombining species. However, the generally clonal and asexual reproduction of *C. albicans* creates a population structure that confounds traditional GWAS methods. By sampling multiple isolates from each individual, we were able to obtain phenotypically diverse strains with a limited set of unique SNPs between isolates, allowing us to identify causative variants associated with a particular phenotype.

We focused on the strains from donor 814, as this donor’s matched oral and fecal samples produced the most colonies of any donor, with 144 colonies arising from the oral sample, and 19 arising from the fecal sample, for a total of 163 colonies. 6 of these 163 colonies were included in the condensed set, and these 6 strains clustered tightly in Clade 1, which we hypothesized would allow us to identify causative variants associated with particular phenotypes that were divergent between strains.

Our agar invasion analysis revealed that isolate 814-168 demonstrated hyper invasion into Spider agar at 37°C, whereas the other 5 isolates from the condensed set were less invasive (Fig 5A). Moreover, from this donor’s 163 isolates, only this single isolate exhibited the hyper invasive phenotype into Spider agar at 37°C (Fig 5B); this phenotype was the motivation for initially including this strain in the sequenced set.

**Fig 5.**
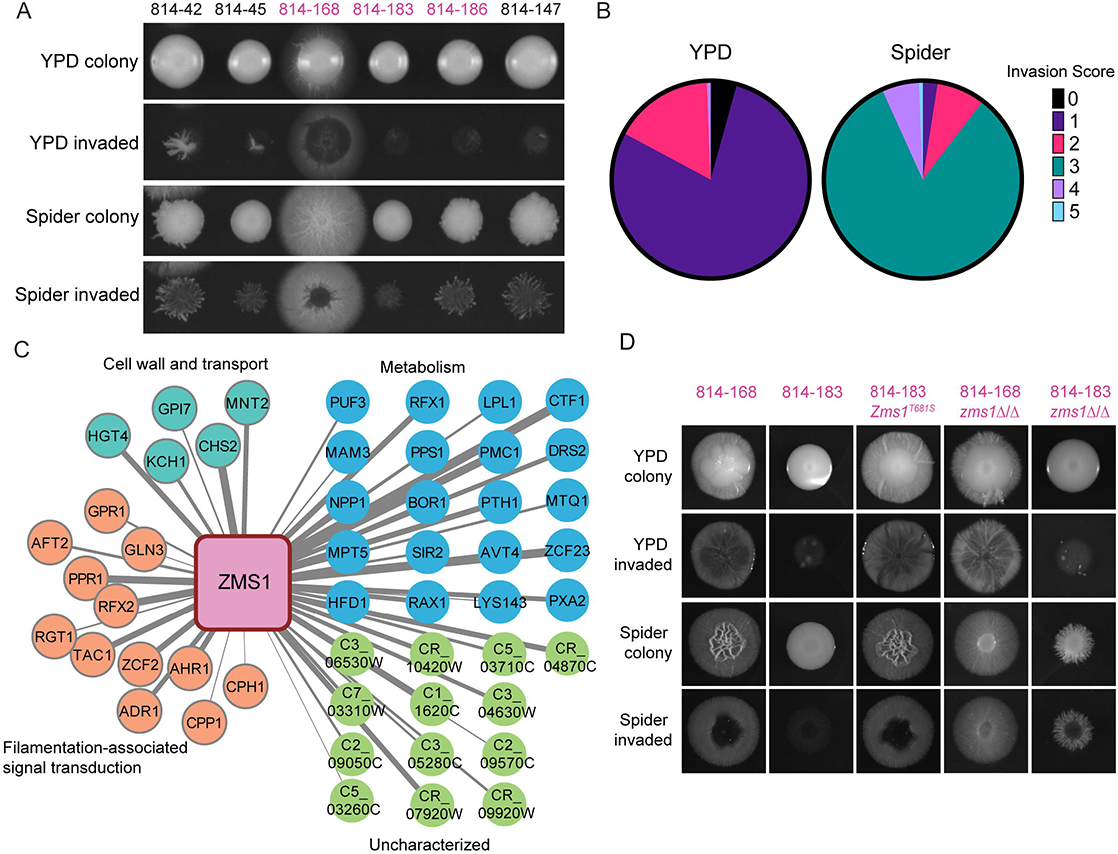
SNV Limited Diversity Exploitation analysis of donor 814 commensal isolates reveals role for Zms1 in regulating agar invasion. A) Agar invasion images for isolates from donor 814 included in the condensed set. Colonies were grown on YPD or Spider agar for 5 days. Invasion was determined after gentle washing. B) Agar invasion scores for all isolates from donor 814 under YPD and Spider conditions. C) Co-expression analysis of Zms1. Width of the lines represents strength of the co-expression score. Gene names and predicted functions from Candida Genome Database. Dark outlines indicate genes examined in Fig 5E. D) Agar invasion images for allele swap and deletion strains. Colonies were grown on YPD or Spider agar for 5 days before imaging. Invasion was determined after gentle washing.

Variant analysis identified 12 genes with unique SNVs in the 814-168 strain compared with the other 5 sequenced 814 strains, including a heterozygous adenine to thymine SNV in the transcription factor Zms1, resulting in a change in amino acid 681 from a threonine to a serine. Moreover, co-expression analysis [64] of *ZMS1* revealed that it is highly correlated with genes involved in regulating the yeast-to-hyphal morphogenic transition (Fig 5C). To test whether this SNV can drive an invasive phenotype, we generated complementation plasmids encoding the *ZMS1T681S* allele cloned out of the 814-168 background, and performed allele swap transformations into 814-183, a closely related strain from the same host which demonstrated minimal agar invasion. In the minimally invasive 814-183 background, replacing one copy of the endogenous *ZMS1* allele with one harboring a serine residue at amino acid 681 (T681S) resulted in hyper invasion into Spider agar (Fig 5D).

Previous work on the function of Zms1 via deletion mutant analysis had not revealed a phenotype [65], however, this was in the SC5314 genetic background and the impact of a specific transcription factor on a given phenotype can vary depending on the strain [17]. Therefore, we deleted *ZMS1* from both 814 backgrounds and tested the strains for invasion and filamentation. On YPD agar, deletion of *ZMS1* in both genetic backgrounds had minimal effects, with the mutant strains behaving similarly to their parent strains (Fig 5D). However, on Spider agar, *ZMS1* deletion changed the colony morphology in the 814-168 background, although it did not decrease overall invasion. Additionally, deletion of *ZMS1* increased invasion in the otherwise minimally invasive 814-183 background, highlighting the differential impact of *ZMS1* mutation in the different genetic backgrounds (Fig 5D). Our results demonstrate that a single SNV changing amino acid 681 to a serine is a dominant active allele that is sufficient to drive a hyphal invasion program into Spider agar. We also identified natural variation that was distinct from deletion phenotypes. This approach highlights how deep phenotypic analysis of a limited set of natural isolates from a single host can be exploited to identify causative variants and identify new functions for under-characterized genes.

### Virulence

We next examined the fitness and virulence of the commensal isolates relative to the SC5314 reference strain; we hypothesized that the commensal isolates would have decreased virulence compared to SC5314, a clinical isolate. To test this, we turned to two models of *Galleria mellonella* systemic infection as this insect model of virulence is significantly correlated with the murine systemic infection model [18,66,67]. We first examined competitive fitness by infecting with a 1:1 mixture of fluorescently marked SC5314 and each sequenced commensal isolate [68]. After three days, the worms were homogenized and CFUs were plated to determine the competitive index, calculated as the log2 ratio of fluorescent to nonfluorescent colonies [35]. When comparing the marked and unmarked SC5314 strains, we obtained a competitive index of 0, indicating that both strains are equally fit and that the fluorescence does not impose a fitness cost. In contrast, most commensal strains had a competitive index > 2, suggesting that these strains have a competitive advantage over SC5314, even during systemic infection (Fig 6A). Notably, the 3 strains previously identified from Clade C, 882-60, 882-46, and 811-7, consistently demonstrated a competitive index of less than 0, indicating these strains are less fit than SC5314 in this model of infection. This is consistent with the slow growth exhibited by these strains in many growth conditions (Fig 4A).

**Fig 6.**
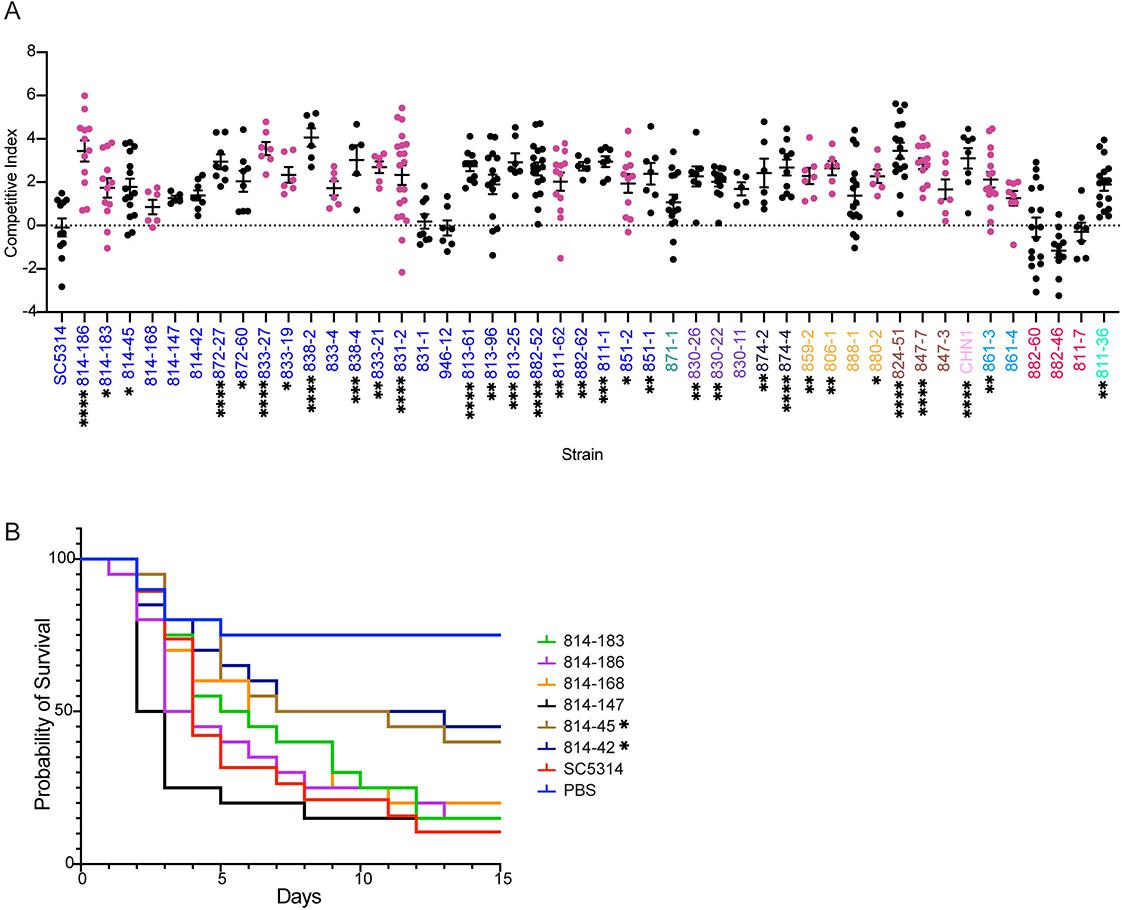
Commensal isolates retain pathogenic potential. A) Competition assays in *G. mellonella* demonstrate increased fitness of many commensal isolates compared to SC5314 reference. Isolates were competed against a fluorescent SC5314 isolate, starting at a 1:1 initial inoculum. Competitive fitness was calculated as the ratio between fluorescent and non-fluorescent colonies, normalized to the inoculum, and log2 transformed. Significant differences from the SC5314 reference strain were determined by one-way ANOVA, with Dunnett’s multiple correction testing. B) Survival assays in *G. mellonella*, comparing the SC5314 reference to 6 isolates from donor 814. Each strain was standardized to 2×10^6^ cells/mL before inoculating 20 *G. mellonella* larvae per strain with 50 µL of prepared inoculum. Larvae were monitored daily for survival. Statistical differences were determined using a Mantel-Cox log-rank test. Asterisks indicate P < 0.05 (*), P < 0.005 (**), P < 0.001 (***) and P < 0.0001 (****) compared with SC5314.

The striking increased competitive fitness of the other isolates motivated us to test whether this increased ratio was correlated with increased disease. Here, we examined the survival of *G. mellonella* after performing monotypic infections. We started by using the six isolates from donor 814, which all had increased competitive fitness compared with SC5314, to test the hypothesis that increased invasion is associated with increased disease. However, the majority of the strains from donor 814 were not significantly different from the SC5314 reference strain (p > 0.05, log-rank test), including the hyper-invasive 814-168 isolate (Fig 6B). Two strains had a slight defect in virulence (p < 0.05 log-rank test). We additionally tested two other clusters of strains for their ability to cause systemic disease in the insect model. However, the human commensal isolates were again not significantly decreased in virulence from the SC5314 reference strain (S6 Fig), despite the increased ability of the SC5314 strain to cause host immune cell death (Fig 4C). Together, this suggests that there is not a consistent selection for avirulent behavior in the commensal human isolates. Although we observed wide variation in *in vitro* host response phenotypes, the isolates generally retained their pathogenic potential, indicating that our *in vitro* assays may not capture the complex stresses experienced during whole-organism infections.

## DISCUSSION

Intra-species analyses of microbial strains can allow for the identification of variants that are associated with specific clinical outcomes, as demonstrated widely in bacterial virulence [69, 70] and recently in the human fungal pathogen *Cryptococcus neoformans* [71, 72]. Understanding the differences between strains can allow for insights into mechanisms of colonization and pathogenesis [17]. However, assessing the underlying genetic variation responsible for differences in virulence in *C. albicans* is challenging in the absence of clear candidate genes because *C. albicans* is a primarily clonal yeast that does not generally undergo meiosis and recombination. Additionally, the genetic basis underlying the ability of commensal strains of *C. albicans* to transition to pathogenic behavior is not well defined. Moreover, *C.albicans* exhibits high rates of structural mutation and ploidy variation [41, 73] as well as high heterozygosity [24]. Previous descriptions of population genomic variation in *C. albicans* have largely relied on short read sequencing [24] or molecular typing methods [27] at relatively few loci to define genetic variation in the species. Here, we have greatly expanded the number of available long-read genomes for *C. albicans* and have generated a catalog of structural variants that was largely absent from previous descriptions of natural diversity. We were able to leverage clonal variation within a single donor to identify meaningful variation that arose during microevolution. Overall, we performed a systematic phenotypic analysis of commensal isolates from healthy donors, thus allowing us to examine *C. albicans* genotypic and phenotypic diversity before the transition to virulence.

Strikingly, our commensal strains generally maintained their capacity to cause disease, and all strains were able to filament in response to the inducing cues of 10% serum or high temperature. This observation is in contrast to work suggesting that passage through the mammalian gut would cause a decrease in systemic virulence and filamentation [34]. Although we observed significant variation in the ability of strains to invade into agar, recent intravital imaging approaches suggest that filamentation in response to serum matched that seen *in vivo* more than filamentation in response to solid Spider agar [74]. Together, the robust filamentation and disease-causing ability in all of the commensal isolates suggests that the selective pressures that occur during mouse models of colonization may not recapitulate the selection that occurs during human colonization.

Although we were not able to associate a particular phenotype with increased systemic disease, we observed extensive phenotypic and genotypic variation between the commensal isolates. This variability is also consistent with recent work from clinical, disease-associated strains [18]. Moreover, our commensal strains were able to proliferate on a range of different environmental conditions, and we did not observe significant differences in phenotypes between isolates obtained from oral or fecal sites, except for a slight increase in invasion into Spider medium for oral samples. This work highlights the striking ability of *C. albicans* to adapt to a wide range of environmental conditions and indicates that colonization at a specific body site does not necessarily predict pathogenic potential.

Previous work has suggested that the *C. albicans* population within a given individual is clonal [68,75–78], and that the fungus is acquired during birth as a part of the normal microbiota [42]. In these studies, they were often sampling from patients with active disease; this may suggest that there is selection for the ability to cause disease, resulting in repeated isolation of representative samples of a clonal population. In contrast, we identified disparate individuals that appeared to be colonized by strains that were nearly identical, suggesting that there was some transmission between individuals. Whether these transmission events allow for long-term colonization, and how they affect the initial *C. albicans* colonizing strains, is still not fully understood.

We also observed diversification within hosts, with multiple instances of closely-related strains showing variation in phenotypes, such as invasion into agar. In one representative example, we were able to use comparative genome analysis coupled with co-expression, which we term “limited diversity exploitation”, to identify a candidate transcription factor that regulates invasion. Previous work on a *ZMS1* knockout strain of *C. albicans* did not show any differences in phenotype compared with the parent strain [65]. However, we observed that a single amino acid substitution in the predicted fungal transcription factor regulatory middle homology region was sufficient to drive hyper-invasive growth. Moreover, this phenotype was distinct from the deletion phenotypes in these two genetic backgrounds, again contrasting with the SC5314 reference strain. It is likely that populations of colonizing *C. albicans* will vary in other important clinically-relevant traits, including filamentation in macrophages or intrinsic drug tolerance and resistance, and our results suggest that studying more than a single representative isolate gives opportunities to discover new biology.

## METHODS

### Human Subjects and Sample Collection

Study volunteers were recruited through the Authentic Research Sections of the introductory biology laboratory course at the University of Michigan (BIO173). All participants provided written, informed consent for sample collection as well as isolation and characterization of microbes from feces. This study was approved by the Institutional Review Board of the University of Michigan Medical School (HUM00094242 and HUM00118951) and was conducted in compliance with the Helsinki Declaration.

Oral samples were obtained by self-administered cotton swabbing, and fecal samples were collected and diluted in PBS+DMSO before plating; in each case, BD ChromAgar was used to identify *C. albicans* colonies. Individual *Candida albicans* colonies were picked from all plates with a toothpick into 100 µL of YPD in a 96 well plate. The cultures were then inoculated onto BD Chromagar to differentiate between yeast species and ITS regions were amplified to confirm *C. albicans* strains. The *C. albicans* colonies were individually inoculated into fresh 96-well plate in YPD and after 24 hours of incubation at 30° C, 50% glycerol was added to generate the stock plates. For each oral sample, individual *Candida albicans* colonies were picked from the CHROMagar plates and incubated overnight in 100 µL of YPD in a 96 well plate. After 24 hours incubation in 30° C, 50% glycerol was added to the 96 wells to generate the stock plates. All strains were maintained in −80 C cryoculture.

### Media and culture conditions

Media for growth curves is described in Supplemental Table 1.

### Growth Curve Analysis

Overnight cultures of each *Candida albicans* isolate were grown in 200 μL of YPD in 96-well plates at 30° C, then subcultured into fresh media using a sterile pinner. Growth curves were performed on a BioTek 800 TS Absorbance Reader in the indicated conditions for 24 hours, without shaking. Maximum growth rate and carrying capacity were determined using the GrowthcurveR analysis package [79]. All media and conditions are listed in Table S1. Summary statistics from GrowthcurveR for the entire collection of isolates are included in Table S2. Summary statistics from GrowthcurveR for the condensed set in multiple environmental conditions are included in Table S4.

### Filamentation Analysis

Overnight cultures of each *Candida albicans* isolate were grown in 100 μL of YPD in 96 well plates at 30° C, then subcultured into 100 μL of YPD at either 30° C or 42° C for three hours. After three hours, the plates were imaged on an Olympus IX80 microscope at 20X magnification. To test the response to serum, overnight cultures were subcultured into 5 mL of YPD with 10% serum and rotated in a 37 °C incubator for three hours before vortexing and imaging on glass slides at 20X magnification.

### Agar Invasion Methods

Isolates were taken from frozen glycerol stocks, incubated overnight in yeast extract peptone dextrose (YPD) in 96-well plates and inoculated onto solid YPD and spider media using a 96-pin replicator tool (Singer Instruments). Plates were then incubated at the indicated temperature and oxygen conditions for 7-9 days. Colonies on plates were first imaged on a Biorad Gel Doc XR. Colonies were then gently washed off with deionized water. Plates were imaged again to capture invasion images. The invasiveness of each isolate was determined through a rubric scale of 0-5, as indicated in Fig 1. A 0 indicates no invasion, 1-2 indicates minimal invasion, a 3 indicates distinct circular hyphal invasion, while a 4 indicates an even larger circular hyphal invasion. A 5 indicates the most aggressive invasion, with a “halo” of more hyphae surrounding the initial growth. A summary of invasion scores is in Table S2.

### Galleria mellonella Infections

Each isolate was examined for competitive fitness in a one-to-one ratio with a fluorescent wildtype SC5314 *C. albicans* strain in *Galleria mellonella* larvae. Infections were performed as previously described (Fuchs et al., 2010). Briefly, *G. mellonella* larvae were purchased from speedyworm.com and maintained in sawdust at room temperature. Overnights were prepared for each isolate and wildtype strain in yeast extract peptone dextrose (YPD) at 30°C, with rotation.

To measure competitive fitness, a 1:1 ratio of fluorescent wild type to unlabeled isolate was prepared in 1X PBS at 5×10^5^ cells/ml. 10 larvae/strain were randomly chosen and infected via the last right proleg with 50 µL of the 1:1 inoculum using an exel veterinary U-40 diabetic syringe (0.5CC X 29G X ½). After injection, larvae were maintained at room temperature for 3 days before harvesting using a Benchmark D1000 homogenizer. Larvae were homogenized in 0.5 ml of 1X PBS, diluted 1:10 in PBS, and plated onto YPD plates containing gentamicin, ampicillin, and ciprofloxacin. Plates were left at 30°C for 48 hours before imaging on a Typhoon™ FLA 9500 biomolecular Imager. The ratio of fluorescent to unmarked strains was compared with the inoculum to determine competitive index.

To measure virulence, 20 *G. mellonella* larvae*/*strain were infected with 50 µL of 2×10^6^ cells/mL inoculum using an exel veterinary U-40 diabetic syringe (0.5CC X 29G X ½). After injection, larvae were maintained at room temperature and monitored daily for survival. Virulence was analyzed using Kaplan Meier survival curves in GraphPad Prism (version 9).

### Mammalian Cell Culture

J2-iBMDM cells were isolated from the bone marrows of C57BL/6J mice and differentiated in BMDM medium (50% DMEM, 2 mM l-glutamine, 1 mM sodium pyruvate, 30% L929-conditioned medium, 20% heat-inactivated fetal bovine serum [FBS; Invitrogen], 55 µM 2-mercaptoethanol, and Pen/Strep), then immortalized using Cre-J2 viral supernatants.

### Phagocytosis

Phagocytosis assays were performed as previously described [58]. Briefly, iBMDM macrophages were prepared for infection in RPMI media containing 3% FBS, diluted to 3×10^6^ cells/ml and incubated overnight at 100 µL/well in a 96-well plate. Overnight cultures of *Candida albicans* isolates were incubated and diluted to 4 × 10^6^ cells/ml into RPMI media with 3% FBS and used to infect macrophages. Inoculated plates were centrifuged for 1 minute at 1,000 rpm to synchronize. After 30 minutes, media was removed, and cells were fixed with 4% paraformaldehyde (PFA) for 10 minutes. The cells were washed three times in 1 X PBS. Cells were then stained with 50 µL of FITC-Concanavalin (5 ug/mL Sigma-Aldrich C7642) at room temperature, rocking for 30 minutes, wrapped in foil. The plates were then washed 3X with 100 µL of 1X PBS, and then 50 µL of 0.05% Triton-X100 was added to permeabilize the cells. Cells were then washed 3X with 1X PBS and a final stain of 50 µL of calcofluor white (100 ug/mL, Sigma-Aldrich C7642) was added to cells to incubate for 10 minutes. The cells were then washed 3X with 100 µL of 1X PBS and then maintained in 100 µL of 1X PBS at 4°C before imaging on the microscope at 20X magnification using the DIC, FITC, and DAPI channels. Images were analyzed with a CellProfiler pipeline to determine the percentage of internalized *Candida*. The total number of *Candida* was determined through calcofluor white staining. Next, the number of external *Candida* was determined through FITC-ConA staining. (1 – external cells)/total cells = % of internalized *Candida*.

### Macrophage filamentation assay

To assess filamentation of isolates in macrophages, iBMDM macrophages were prepared for infection in RPMI media containing 3% FBS, diluted to 3×10^6^ cells/ml and incubated overnight at 100 µL/well in a 96-well plate. Overnights of *Candida albicans* isolates were incubated and diluted to 4 × 10^6^ cells/ml into RPMI media with 3% FBS and used to infect macrophages at an MOI of ∼1:1. The inoculated plate was centrifuged for 1 minute at 1,000 rpm to synchronize phagocytosis. After 2 hours, the media was removed, and cells were fixed with 4% paraformaldehyde (PFA) for 10 minutes, permeabilized with 0.05% Triton-X100, and stained with 50 µL of calcofluor white (100 ug/mL, Sigma-Aldrich C7642) before imaging at 40X magnification using the DIC and DAPI channels. The extent of filamentation for each isolate was scored on a rubric from 0-5. A score of 0 indicates an isolate that exhibited only yeast morphology with no filamentation during macrophage infection. A score of 1 indicates an isolate that remains primarily in the yeast form, with some hyphae or pseudohyphae, while a score of 2 indicates an isolate that remains primarily in the yeast form, with more hyphae and pseudohyphae present than a score of 1. A score of 3 indicates an isolate that had approximately equal numbers of yeast and true hyphae. A score of 4 indicates an isolate with more true hyphae than yeast during infection and a score of 5 indicates an isolate in which the majority of cells formed hyphae with very few yeast remaining.

### Cell death assay

To assess macrophage cell death, J2-iBMDM macrophages were prepared for infection in RPMI media containing 3% FBS, diluted to 4×10^5^ cells/ml and incubated overnight at 100 µL/well in a 96-well plate. Macrophages were primed with 200 ng/mL LPS in RPMI media with 3% FBS for 2 hours. Overnights of *Candida albicans* isolates were incubated and diluted to 5 × 10^5^ cells/ml into RPMI media with 3% FBS and used to infect macrophages at an MOI of ∼1:1. The plate was centrifuged for 1 minute at 1,000 rpm to synchronize phagocytosis. After 4 hours, the media was removed and cells were stained with Hoechst (20 uM, Cayman Chemical 33342) to mark nuclei and propidium iodide (1ug/mL, Acros Organics 440300250) to mark cell death. Plates were then imaged on an Cellomics ArrayScan™ VTI Objective Module at 20x with at least 5 images taken per well. Images were analyzed with a CellProfiler pipeline (available by request) to determine the percent of dead macrophages. Macrophages were identified through Hoescht nuclear staining to determine the total number of cells in each field of view. Next, cell death events were determined based on propidium iodide staining. (# of propidium iodide events)/(# of total cells) = % of dead macrophages.

### Biofilm Formation

Biofilm formation was assessed as previously described [56]. Briefly, overnights of *C. albicans* isolates were incubated and diluted to 0.5 OD_600_. In a 96-well plate, isolates were added to 200 μL of RPMI media, covered with a Breathe-Easy film, and incubated at 37°C shaking at 250 rpm. After 90 minutes, RPMI media was removed and wells were washed once with 200 μL of 1X PBS and 200 μL of fresh RPMI media was added. Plates were covered in a Breathe-Easy film and left to incubate at 37°C with shaking at 250 rpm for 24 hours before a final read on the plate reader at OD_600_.

### CHEF gel electrophoresis

Agarose plugs for CHEF gel electrophoresis were prepared as in Selmecki 2005 [80]. Electrophoresis was performed as in Chibana [81], with minor modifications to improve chromosome separation. Plugs were run on 0.9% megabase agarose (Biorad 1613108), with run conditions of 60 to 300 sec, 4.5 V/cm, 120 angle for 36 hr, followed by 720 to 900 sec, 2.0 V/cm, 106 for 24 hr. Sizes were determined based on approximations from Selmecki and a CHEF DNA Size Marker, 0.2–2.2 Mb, *S. cerevisiae* Ladder (Biorad 1703605).

### High Molecular Weight Genomic DNA extraction and sequencing

High molecular weight (>20 kb fragment) genomic DNA was prepared using the Qiagen MagAttract HMW DNA Kit (Qiagen 67563), following the manufacturer’s protocol, with minor modification. Fungal cell walls were first digested using Zymolyase (Zymo E1005) before DNA extraction. TELL-seq [48] libraries were prepared at the University of Michigan Advanced Genomics facility. Libraries were sequenced on an Illumina NovaSeq SP 300 cycle flow cell. All sequences are uploaded to NCBI SRA at PRJNA875200.

### Sequence Variation Analysis

To identify single nucleotide variants (SNVs), we mapped 435 read libraries to the SC5314 reference genome with BWA-MEM [49]. We used various samtools utilities to convert alignment files, remove PCR duplicates, and assign read groups [49]. Variants were called with GATK HaplotypeCaller and individual sample GVCFs were combined with GATK CombineGVCFs. Finally, we genotyped our combined GVCF across samples and loci with GATK GenotypeGVCFs [82]. GATK identified 851,355 total variable loci. Of the 741,027 diallelic loci, GATK identified 60,626 indel-derived alleles (8.18%), and 680,401 SNV alleles (91.82%). To refine our raw variant set to only include high-confidence SNVs, we removed low quality variant calls at the (i) individual genotype call, (ii) whole locus, and (iii) whole sample levels. First, we removed individual sample genotype calls if their genotype quality (i.e., GQ) was < 0.99 (out of 1.0) or the results of the map quality rank sum test (i.e., MQRS) differed from 0.0 (i.e., the perfect score). Second, we removed all variant calls with more than one alternative allele (i.e., not diallelic), an alternate allele longer than 1 nucleotide (i.e., indels), or with >5% missing genotype calls across samples. Finally, we removed entire samples from the combined VCF if their genotype calls across all loci were more than 90% missing, which resulted in the removal of 4 entire samples. Filtering our variant set in this way yielded a final set of 112,136 high quality SNVs across 431 remaining samples. Of these, 90,675 (80.86%) were represented in at least one member of our set of newly sequenced strains.

To remove redundancy and focus on natural *C. albicans* diversification, we removed samples corresponding to resequenced strains (e.g., multiple SC5314 samples present in full data set) and those sequenced as part of experimental evolution studies (i.e., [34,51,52]). Following removal of these samples, we were left with 324 sample SNV profiles (280 published, and 44 new genomes strains) (Table S3). To generate an input matrix for distance calculation and NJ clustering, we coded genotypes as homozygous for the reference allele (R), heterozygous (H), homozygous for the alternative allele (A), or missing (N). Raw distance between two sample genotypes at a particular locus (i.e, GT_i_ and GT_j_) were calculated as 0.5 per alternative allele, with respect to the SC5314 reference such that D(H, A) = 0.5, D(R, A) = 1.0, etc. Missing genotypes (i.e., coded as N) were ignored. Using these raw distances, we calculated the pairwise distance between all sample pairs across all 112,136 loci according to the generalized distance function for sample *i* and sample *j*…

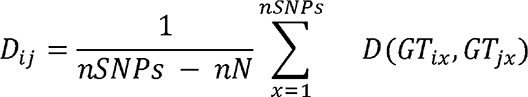

…where *nSNPs* is the total number of high-quality SNPs in our dataset (i.e., 112,136), *nN* is the number of uncalled genotypes between each pair (i.e., coded as N), *GT_ix_* is the genotype of sample *i* at locus *x*, and *GT_jx_* is genotype of sample *j* at locus *x*. The resulting distance matrix was clustered with the *nj* function in phytools in R [83]. We visualized the resulting tree structure and associated data with ggtree in R [84–86].

### Structural Variant Analysis from TELL-seq

To identify structural variation genome wide in the condensed set of *C. albicans* isolates, we mapped the read libraries to the SC5314 reference genome with BWA-MEM [49]. We visualized the read depth across the genome by breaking the reference genome into 500bp bins. Each read was assigned to a single bin based on the first coordinate in the reference genome to which the read aligned. The read counts were normalized across the samples by the average read coverage for each sample. The value for each bin (v_b,s_) for each sample was calculated using the following formula:

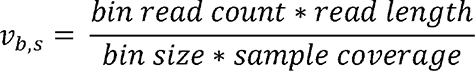

Additionally, we performed structural variant calling using lumpy-sv [87] and genotyping with svtyper [88] using smoove (version 0.2.5). We visualized the variant calls by breaking the reference genome into 500bp bins and identified the number of variant calls of a give type which overlapped with each bin.

To identify potential translocations from the TELL-seq data, we used the Tell-Link pipeline to assemble the read libraries into contigs and aligned the assembled contigs to the SC5314 reference genome using MiniMap2 [89]. We identified candidate junctions where consecutive aligned segments longer than 10,000 bp on the same contig aligned to different chromosomes. To experimentally validate the predicted junctions, we designed primers from uniquely mapped contig segments on either side of the junction i.e., one from each of the two chromosomes predicted to be joined together. Primers were selected from the 500 bp directly upstream and downstream from the predicted junction location unless the junction was directly flanked by non-uniquely mapping segments in which case, the 500 bp of the closest uniquely mapping segments were used. Primers are in Table S5.

### ZMS1 strain construction

To make the *ZMS1* allele swap strains, we used the NEBuilder HiFi assembly kit to clone both alleles of *ZMS1* from the 814-168 strain background into the pUC19 cloning vector, along with the nourseothricin resistance cassette, to generate plasmid pTO192. *ZMS1* and 500 bp of putative terminator was amplified using oTO771+oTO734. NAT was amplified using oTO18 + oTO735. pUC19 was amplified using oTO590 + oTO591. The plasmids were Sanger sequenced to confirm the presence of the specific *ZMS1* allele and the absence of additional mutations using oTO736. These plasmids were transformed into the 814-183 background using a transient CRISPR approach [90]. NAT-resistant transformants were tested for the presence of specific *ZMS1* alleles by Sanger sequencing using oTO736.

To generate the deletion strains, the *ZMS1::NAT* cassette was amplified from the NAT flipper plasmid [91] using primers oTO1215 and oTO1216, and transformed into the 814-168 and 814-183 strain backgrounds using a transient CRISPR approach [90]. Integration was tested using oTO5 and oTO1218 and loss of the wild-type *ZMS1* gene was tested using oTO736 and oTO1218.

## ACKNOWLEDGEMENTS

We thank members of the O’Meara lab for critical reading of the manuscript, the Advanced Genomics Core at the University of Michigan for assistance with TellSeq, and the University of Michigan Great Lakes research computing cluster.

## Supplemental Tables

Supplemental table 1: Media and growth conditions.

Supplemental table 2: Full collection growth curves (Tab 1) and invasion scores (Tab 2).

Supplemental table 3: SRA accession numbers for strains included in phylogenetic analysis.

Supplemental table 4: Condensed set strain information and quantitative phenotypes.

Supplemental table 5: Primers used in this study.

## Supplemental Figure Legends

**Supplemental Figure 1: Comparisons of growth rate between samples and nutrient source.** A) Enrichment of fecal vs. oral samples by carrying capacity. B) Correlation between growth rates in different carbon sources. C) Comparison between oral and fecal strains for agar invasion.

**Supplemental Figure 2: Structural Variation.**

A. CHEF karyotype gels were performed on the condensed set of *C. albicans* isolates and visualized using ethidium bromide staining. Red arrows indicate large chromosome banding patterns that differ from the SC5314 reference strain. Chromosome 5 showed especially extensive variability in size between isolates.
B. Heatmaps of Lumpy Structural variation calls for the condensed set of isolates. Each column represents a 500 bp bin of the reference genome and each row of the heat map for a given variant class is an isolate from the condensed set. The value for each bin indicates the number of SV calls overlapping the 500 bp window.

**Supplemental Figure 2: Biofilm formation.** Each condensed set isolate was tested for its ability to form a biofilm on a plastic surface. Asterisks indicate P < 0.05 (*) compared with SC5314, one-way ANOVA compared with SC5314, with Dunnett’s post-hoc test for multiple corrections.

**Supplemental Figure 4: Comparison between filamentation in macrophages and agar invasion.** A) Images of *C. albicans* condensed set isolates after growth in macrophages for 4 hours. *C. albicans* were stained with calcofluor white. Images taken using DIC and DAPI channels at 20X magnification. Scale = 50 μM. B) Colony morphology and agar invasion for *C. albicans* condensed set isolates on Spider at 37°C after 5 days. C) Histogram of the distribution of macrophage filamentation scores of the condensed set.

**Supplemental Figure 5: UMAP embedding.**

Non-linear embedding dimensionality reduction was performed on the phenotypic data on the condensed set of isolates using UMAP. Clusters did not segregate by sample site, donor, or clade. Data used to generate the UMAP are included in Table S4.

**Supplemental Figure 6: Virulence in *G. mellonella.*** A) Survival assays in *G. mellonella*, comparing the SC5314 reference to 4 isolates from donors 882 and 811. B) Survival assays in *G. mellonella*, comparing the SC5314 reference to 5 isolates from donors 833 and 838. Each strain was standardized to 2×10^6^ cells/mL before inoculating 20 *G. mellonella* larvae per strain with 50 µL of prepared inoculum. Larvae were monitored daily for survival. Statistical differences were determined using a Mantel-Cox log-rank test. ** indicates P-value < 0.01, * indicates P-value < 0.05.

