## Supplemental Figures for "Human commensal *Candida albicans* strains demonstrate substantial within-host diversity and retained pathogenic potential"

Supplemental figure 1

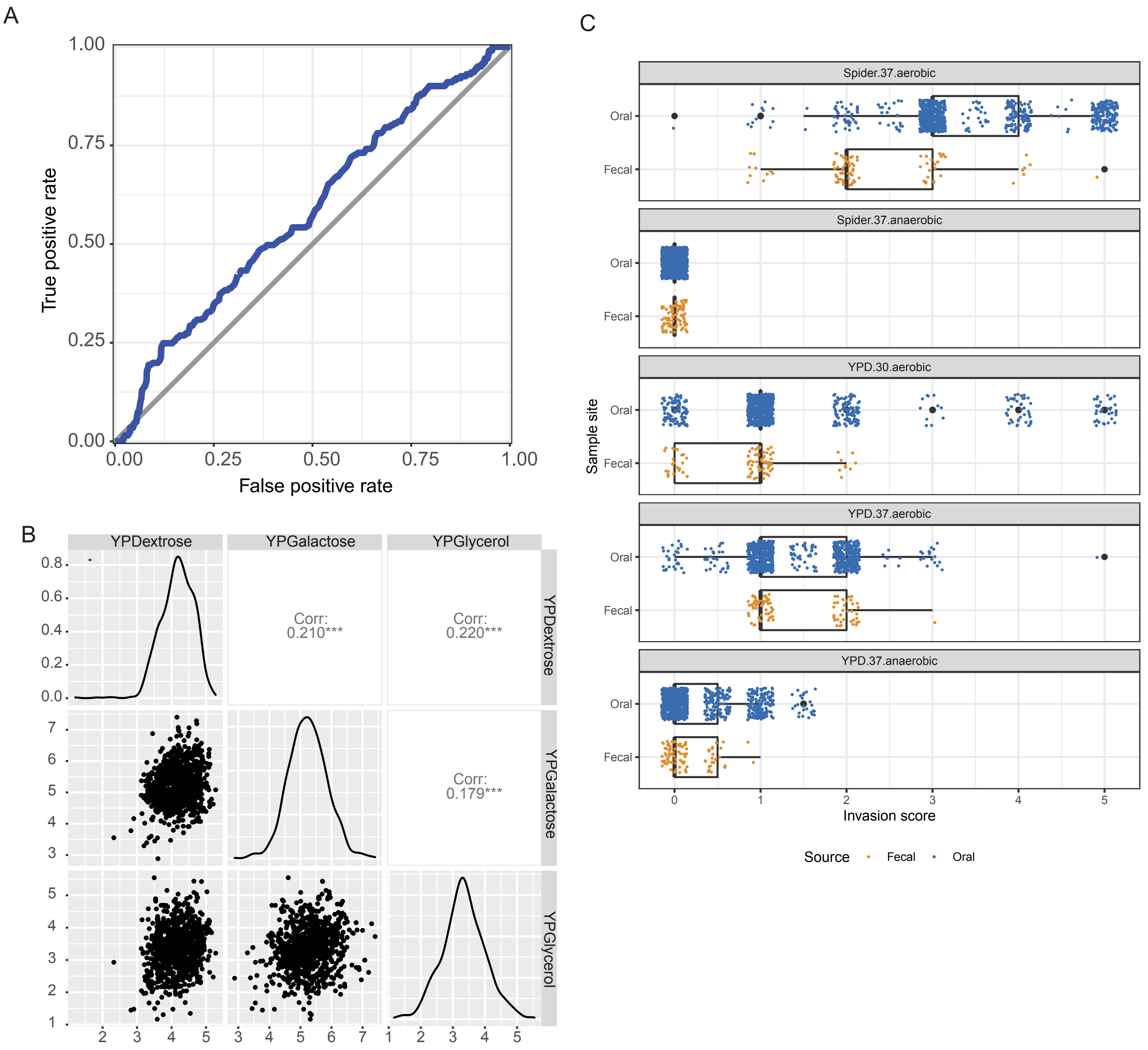

Supplemental figure 2

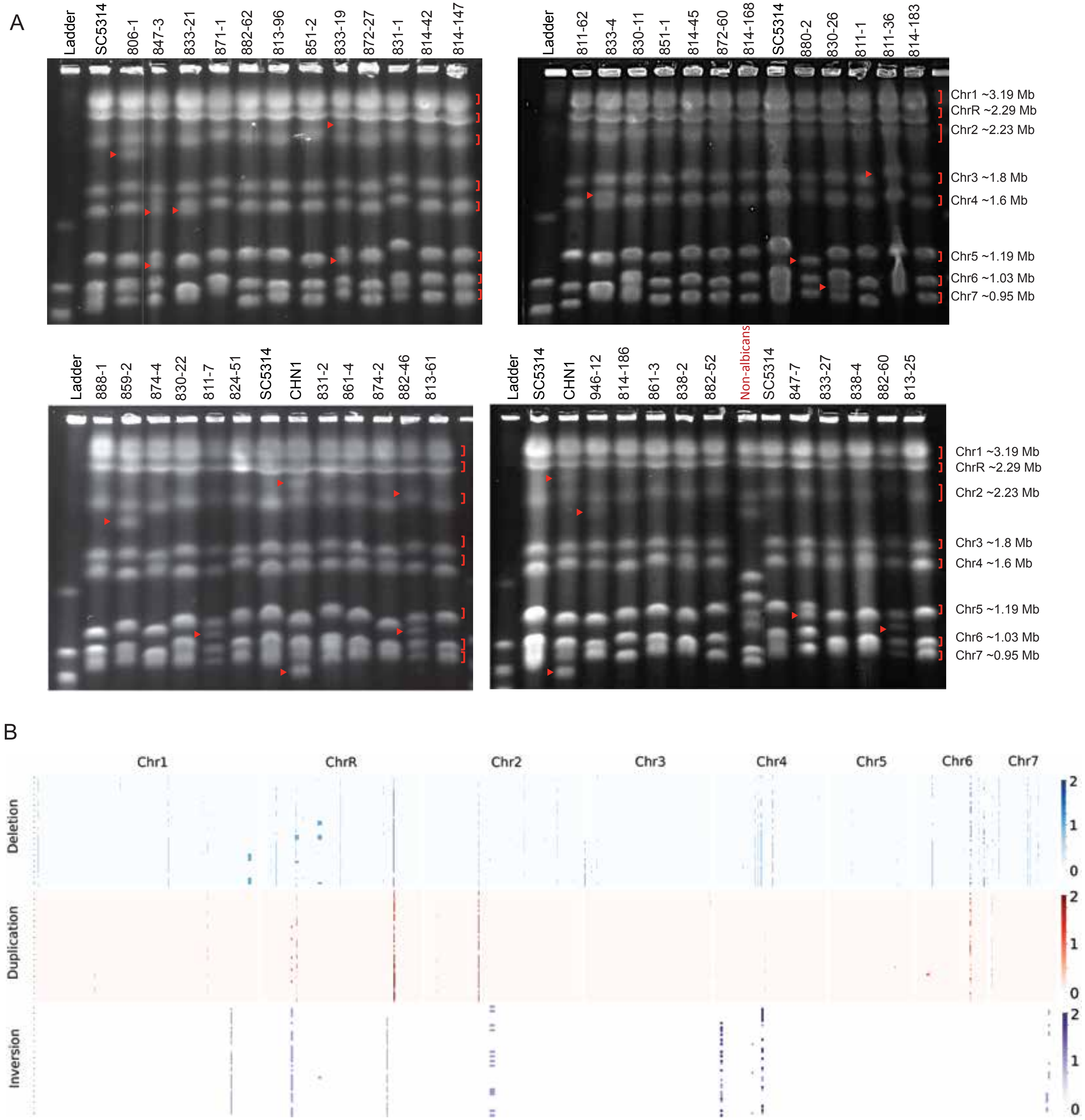

Supplemental figure 3

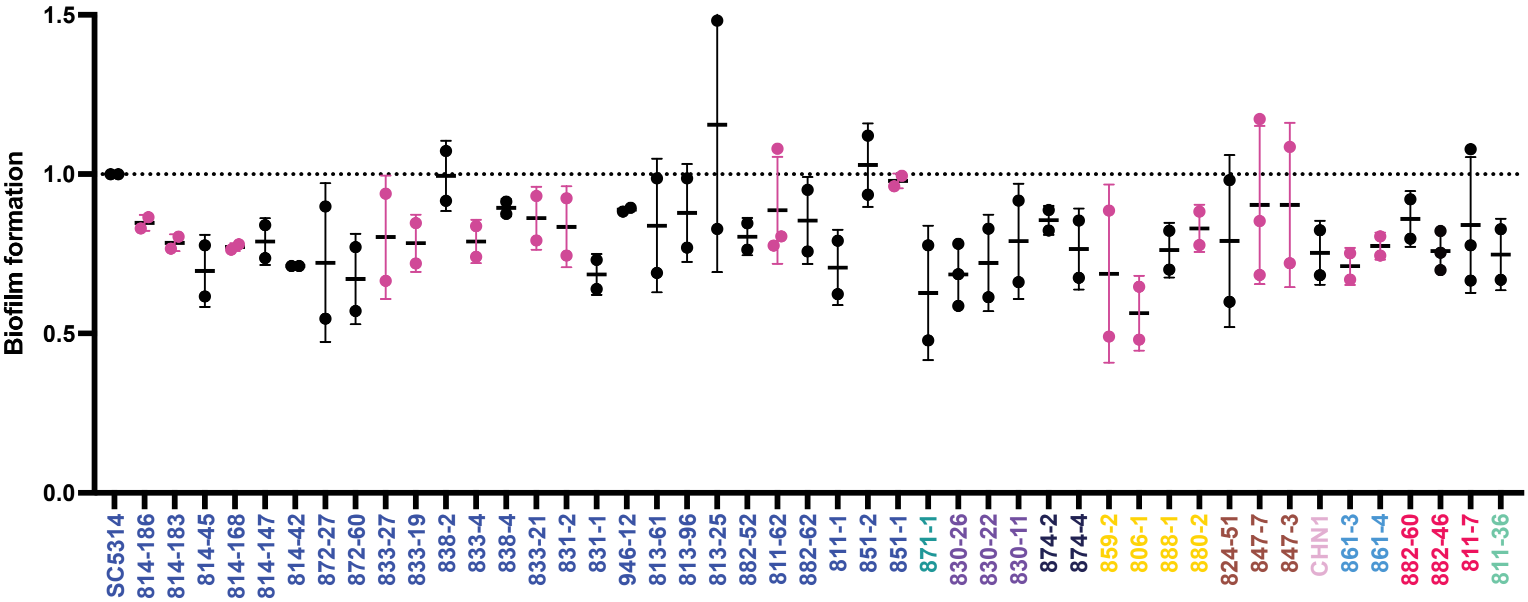

Supplemental figure 4

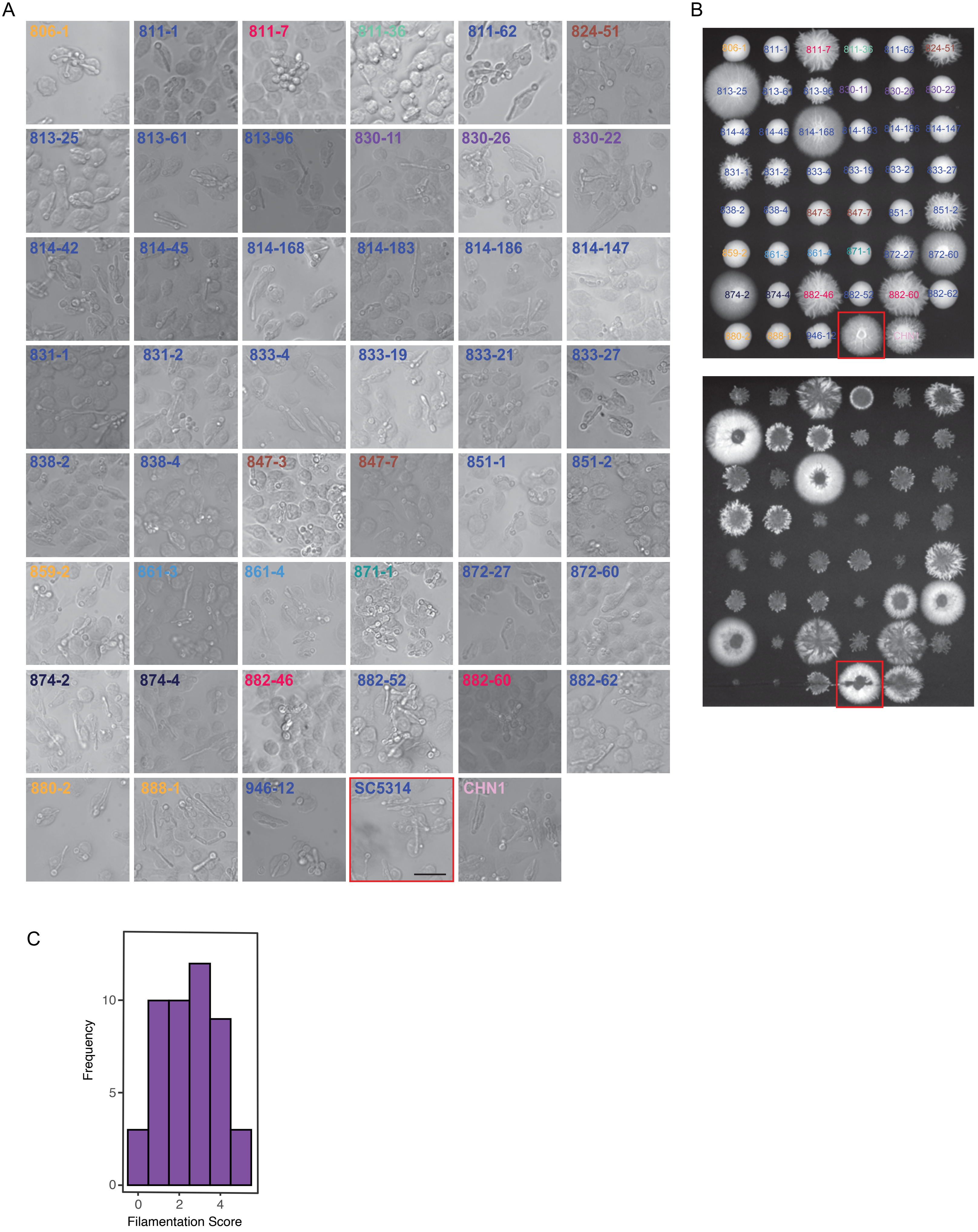

Supplemental figure 5

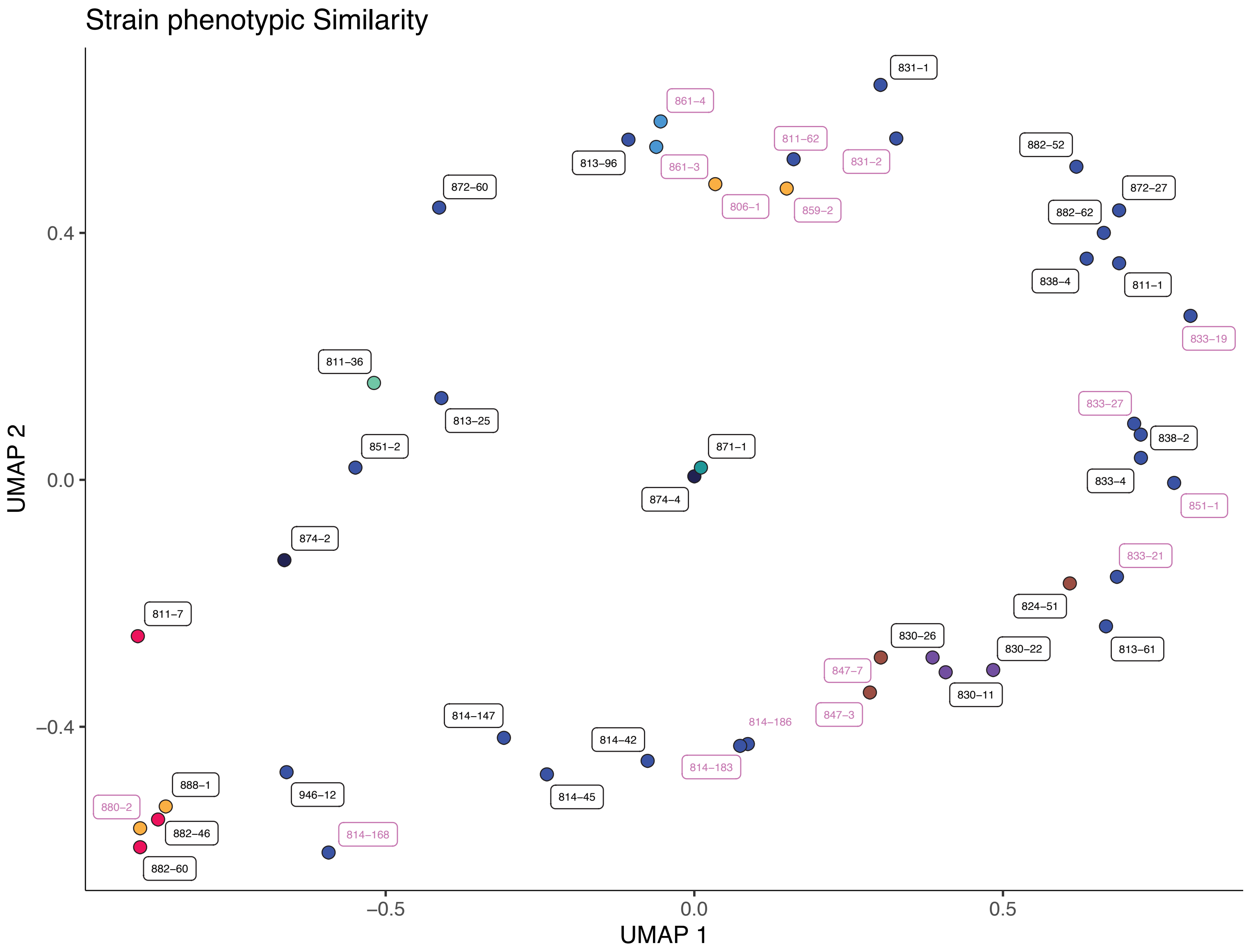

Supplemental figure 6

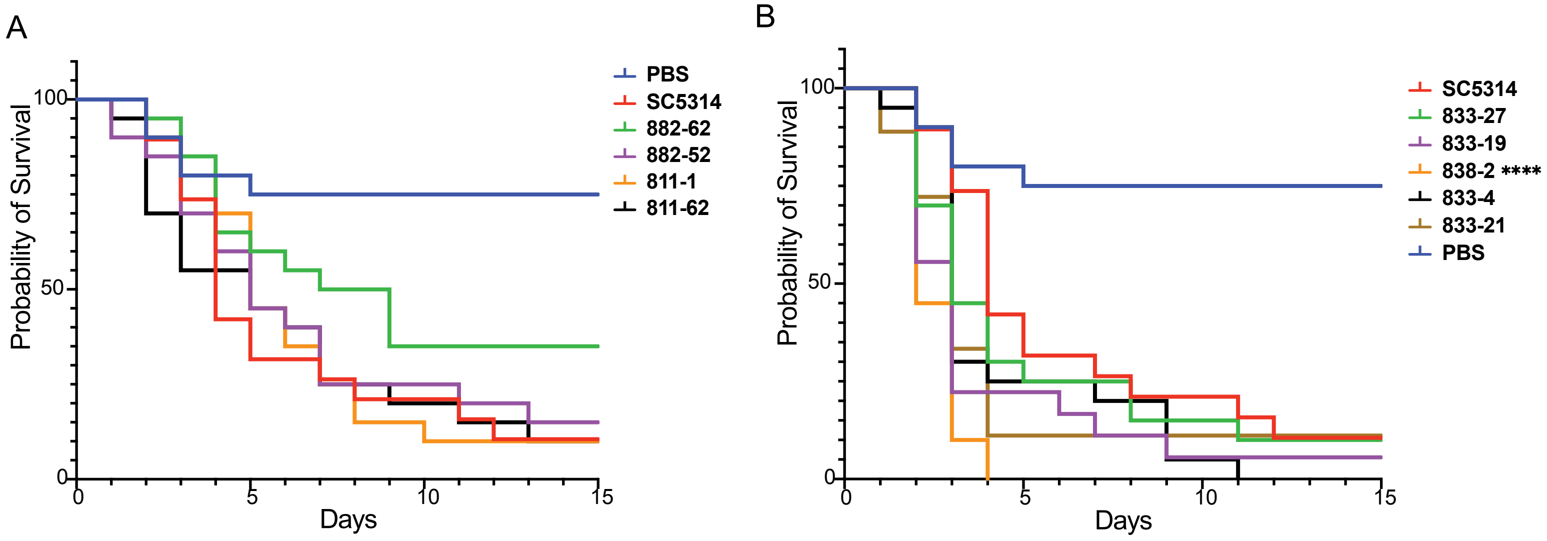
